# The transmembrane proteins M6 and Anakonda cooperate to initiate tricellular junction assembly in epithelia of *Drosophila*

**DOI:** 10.1101/2020.05.01.071746

**Authors:** Anna Wittek, Manuel Hollmann, Raphael Schleutker, Stefan Luschnig

## Abstract

Cell vertices in epithelia comprise specialized tricellular junctions (TCJs) that seal the paracellular space between three adjoining cells [1, 2]. Although TCJs play fundamental roles in tissue homeostasis, pathogen defense, and in sensing tension and cell shape [3-5], how they are assembled, maintained and remodeled is poorly understood. In *Drosophila* the transmembrane proteins Anakonda (Aka [6]) and Gliotactin (Gli [7]) are TCJ components essential for epithelial barrier formation. Additionally, the conserved four-transmembrane-domain protein M6, the only myelin proteolipid protein (PLP) family member in *Drosophila*, localizes to TCJs [8, 9]. PLPs associate with cholesterol-rich membrane domains and induce filopodia formation [10, 11] and membrane curvature [12], and *Drosophila M6* acts as a tumor suppressor [8], but its role in TCJ formation remained unknown. Here we show that M6 is essential for the assembly of tricellular, but not bicellular occluding junctions, and for barrier function in embryonic epithelia. M6 and Aka localize to TCJs in a mutually dependent manner and are jointly required for TCJ localization of Gli, whereas Aka and M6 localize to TCJs independently of Gli. Aka acts instructively and is sufficient to direct M6 to cell vertices in the absence of septate junctions, while M6 is required permissively to maintain Aka at TCJs. Furthermore, M6 and Aka are mutually dependent for their accumulation in a low-mobility pool at TCJs. These findings suggest a hierarchical model for TCJ assembly, where Aka and M6 promote TCJ formation through synergistic interactions that demarcate a distinct plasma membrane microdomain at cell vertices.

## Results and Discussion

### M6 co-localizes with Aka and Gli at tricellular junctions in embryonic epithelia

We analyzed the distribution of M6 in embryos using a GFP protein trap insertion into the *M6* locus, yielding GFP-tagged variants of five of the six annotated M6 isoforms (M6^CA06602^, referred to as GFP::M6; Fig. 1A [13]), or using an antiserum raised against two peptides present in five M6 protein isoforms (Fig. S1A). Anti-M6 immunostaining detected M6 protein in epithelia, including epidermis, salivary glands, tracheae, hindgut, midgut, Malpighian tubules, as well as in the nervous system of stage 14 embryos (Fig. S1D-I). In the epidermis of stage 15 embryos, M6 was detectable at lateral cell membranes (Fig. S1B,C). Compared to anti-M6 staining, GFP::M6^CA06602^ displayed stronger accumulation at TCJs (Fig. S1B), suggesting that anti-M6 preferentially detects M6 isoforms or trafficking intermediates that are not GFP-positive in *GFP::M6^CA06602^* embryos. GFP::M6 accumulated at vertices at the level of septate junctions (SJs) marked by Macroglobulin-/Complement-related (Mcr [14]; Fig. 1B), and co-localized with the TCJ components Aka and Gli (Fig. 1C). In living embryos GFP::M6 was enriched 7.56 +/- 0.66-fold (n=10) at vertices relative to bicellular contacts, comparable to the enrichment of Aka::GFP (6.93 +/- 1.63-fold, n=10) and Gli::YFP (8.61 +/- 1.27-fold, n=10; Fig. 1D-G). These findings suggest that M6 is involved in the formation of TCJs together with Aka and Gli.

**Figure 1.**
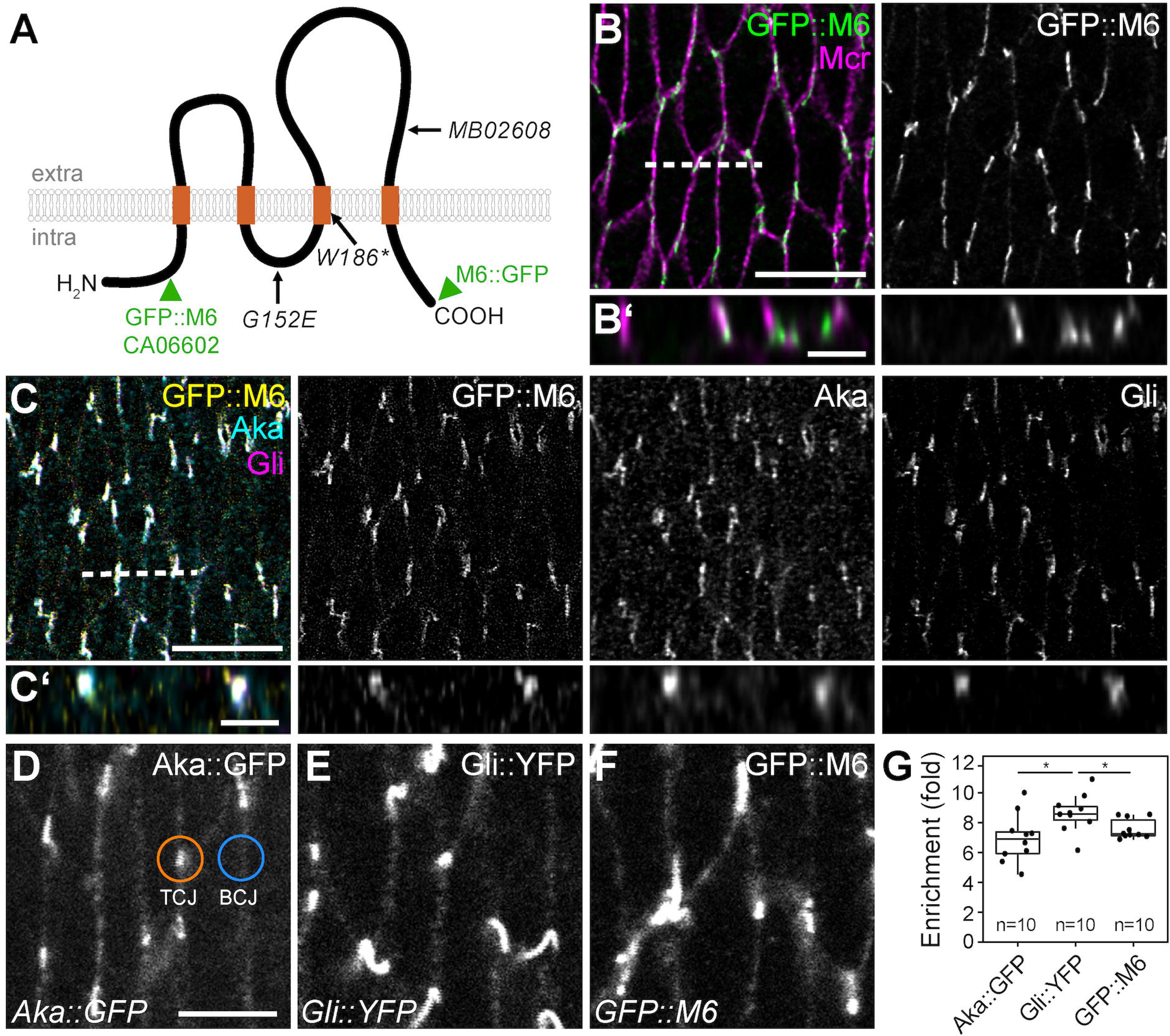
M6 co-localizes with Aka and Gli at tricellular junctions. (A) Schematic drawing of M6 protein with EMS-induced mutations (*G152E, W186\**), Minos transposon (*MB02608*) insertion, and positions of GFP in the CA06602 protein trap line and the UAS-M6::GFP construct (green arrowheads). (B,C) En-face view (B,C) and cross-sections (B’,C’; taken at the dashed line in B,C) of epidermis in stage 16 embryos expressing GFP::M6 (green in B, yellow in C) and immunostained with anti-Mcr (magenta; B) or anti-Gli (magenta; C) and anti-Aka (cyan) (C). GFP::M6 is enriched at TCJs in the basolateral membrane (B) and co-localizes with Aka and Gli (C). (D-F) En-face views of epidermis in living stage 15 embryos expressing endogenously tagged Aka::GFP (D), Gli::YFP (E) or GFP::M6 (F). Note that Aka::GFP, Gli::YFP and GFP::M6 accumulate at vertices and are present at lower concentrations at bicellular contacts. (G) Quantification of enrichment at TCJs measured as the ratio of intensities at tricellular (TCJ) and bicellular (BCJ) contacts (example ROIs shown in D). Sample size (n) is indicated. Wilcoxon rank-sum test, *: p < 0.05. Scale bars: (B,C), 10 μm; (B’,C’), 2 μm; (D-F), 5 μm.

### M6 is essential for epithelial barrier function

To address the role of M6 in TCJ assembly we examined embryos carrying *M6* mutations induced by ethyl methanesulfonate (*W186*, G152E, NC*; [8]) or by a *Minos* transposon insertion (*M6^MB02608^* [15]). M6 protein was detectable by anti-M6 immunostaining in wild-type embryos (Fig. 2H), but not in *M6^MB02608^*, *M6^W186*^* and *M6^NC^* homozygotes (Fig. 2K and data not shown). All four mutants were embryonic lethal as homozygotes or in *trans* to a chromosomal deficiency (*Df(3L)BSC419*) uncovering the *M6* locus. The mutant embryos showed over-elongated tracheae (Fig. 2A,B,E), resembling the defects associated with loss of epithelial barrier function in SJ or TCJ protein mutants [6, 14, 16]. To probe barrier function, we injected Rhodamine-labeled dextran (10 kDa) into the hemocoel of stage 16 *M6^MB02608^* and control embryos. While dextran was excluded from the tracheal lumen in controls (Fig. 2C; n=7), dextran was detectable inside the tracheal lumen in *M6^MB02608^* embryos 20 minutes after injection (Fig. 2D; n=12), indicating a defective epithelial barrier. Thus, *M6* is essential for epithelial barrier function during embryogenesis.

**Figure 2.**
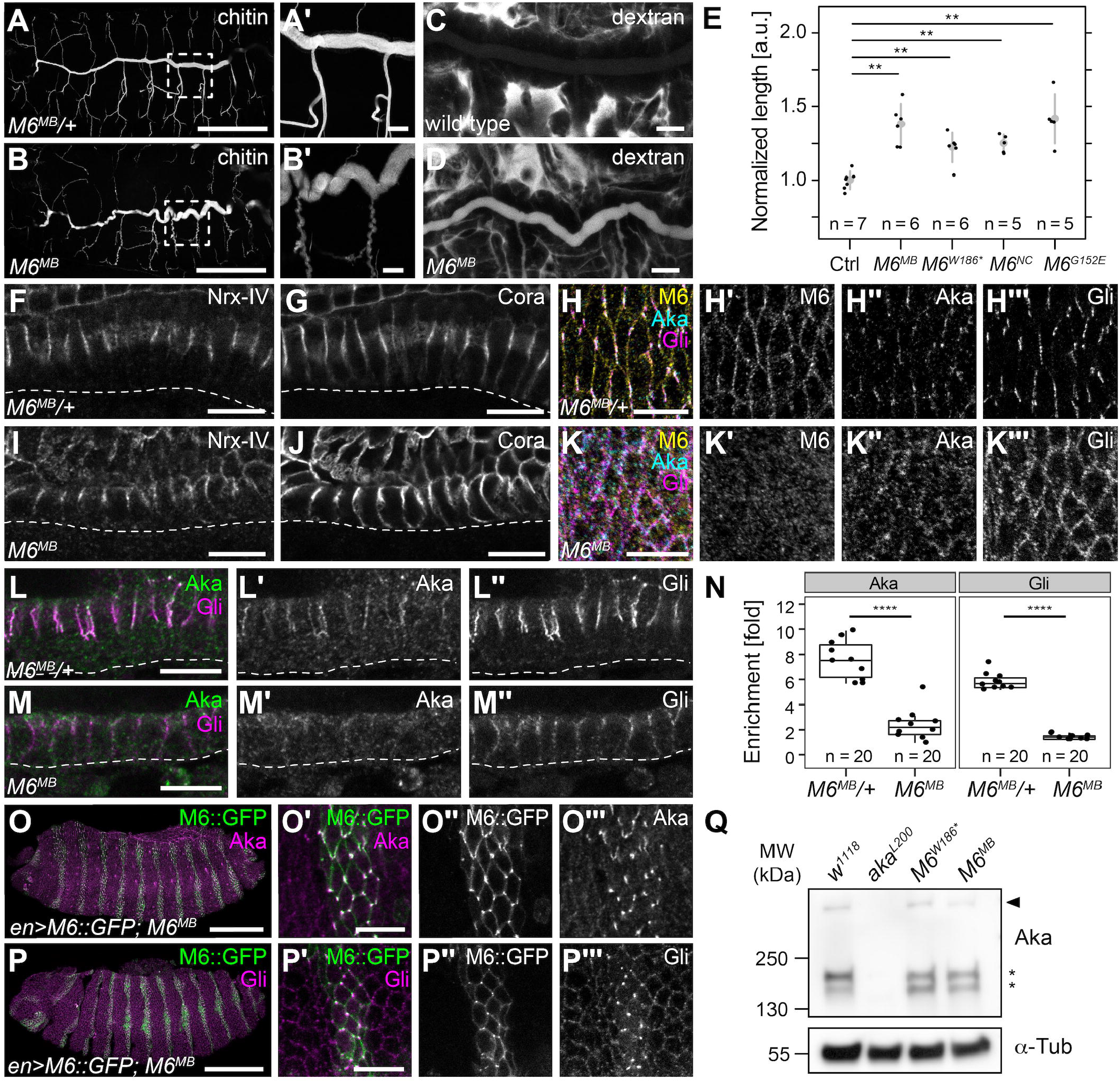
M6 is required for tracheal tube size control, epithelial barrier formation and tricellular junction formation. (A,B) Z-projections of heterozygous *M6^MB02608^/+* (control; A) or homozygous *M6^MB02608^* (*M6^MB^*; B) embryos (stage 16) stained for chitin to label tracheal lumen. (A’) and (B’) are close-ups of regions marked by rectangles in (A) and (B). (C,D) Living embryos (stage 16) were injected with Rhodamine-dextran (10 kDa) and imaged after 20 minutes. Dextran is excluded from the tracheal lumen in wild-type embryos (C), but enters the lumen in *M6^MB02608^* embryos (D). (E) Quantification of tracheal dorsal trunk length (metameres 2-8) in control (*Df(3L)BSC419/+*; Ctrl) and hemizygous embryos carrying the indicated *M6* alleles in trans to *Df(3L)BSC419.* Data is represented as mean +/- S.D. Sample size (n) is indicated. Wilcoxon rank-sum test, **: p< 0.01. (F,G,I,J) Confocal sections of hindgut epithelium in *M6^MB02608^/+* (F,G) or *M6^MB02608^* (I,J) embryos (stage 15) stained for Nrx-IV (F,I) or Cora (G,J). Note that Nrx-IV and Cora *M6^MB02608^* accumulate at apico-lateral membranes in *M6^MB02608^* embryos comparable to controls. Dashed lines mark basal surface of the epithelium. (H,K,L,M) TCJ proteins are mislocalized in *M6* mutants. Control *M6^MB02608^/+* (H,L) and *M6^MB02608^* stage 15 embryos (K,M) were stained for M6, Aka and Gli (H,K) or for Aka and Gli (L,M). En-face view of epidermis (H,K) shows that M6 is detectable at lateral membranes in the control (H’), but is lost in *M6^MB02608^* embryos (K’). Accumulation of Aka and Gli at TCJs in controls (H’’,H’’’) is lost in *M6^MB02608^* embryos (K’’,K’’’). (L,M) Lateral distribution of Aka and Gli in cross-sections of hindgut epithelium in control (L) and *M6^MB02608^* (M) embryos. (N) Quantification of enrichment of Aka or Gli at epidermal TCJs in control and *M6^MB02608^* embryos (stage 15). Sample size (n) is indicated. Wilcoxon rank-sum test, ****: p < 0.0001. (O,P) Expression of *UAS-M6::GFP* rescues TCJ localization of Aka and Gli. *M6^MB02608^*embryos (stage 15) expressing *M6::GFP* in epidermal stripes (driven by *en*-Gal4) were stained for Aka (O) or Gli (P). Note that both proteins are mislocalized in cells between the *en*-Gal4-expressing stripes, but accumulate at TCJs in cells expressing *M6::GFP*. (Q) Immunoblot of extracts from control (*w^1118^*), *aka^L200^, M6^W186^* and *M6^MB02608^* embryos. Positions of full-length Aka (arrowhead) and shorter Aka bands (asterisks) and molecular weight standard (kDa) are indicated. Scale bars: (A,C,O,P), 100 μm; (all other panels), 10 μm.

### M6 is required for assembly of tricellular but not bicellular septate junctions

To understand the role of M6 in barrier function we examined the distribution of two SJ core components, the transmembrane protein Neurexin-IV (Nrx-IV [17]) and the 4.1/Ezrin/Radixin/Moesin (FERM) domain protein Coracle (Cora [18]), in *M6^MB02608^* embryos. Nrx-IV and Cora accumulated at apico-lateral membranes in the hindgut epithelium in *M6* embryos (Fig. 2I,J), resembling their localization in wild-type embryos (Fig. 2F,G), suggesting that M6 is not required for assembly of bicellular SJs. By contrast, the TCJ proteins Aka and Gli were mislocalized in *M6* mutants (Fig. 2H,K,L-N). Instead of accumulating at vertices, Gli distributed evenly around the cell perimeter (Fig. 2H”’,K”’), although it remained enriched at the apico-lateral SJ domain (Fig. 2L’’,M’’). Similarly, Aka accumulation at TCJs was lost in *M6* embryos (Fig. 2H’’,K’’,L’,M’, Fig. S2). In living embryos expressing endogenously tagged Aka::GFP in the absence of M6, Aka::GFP was detectable around the lateral membrane (Fig. S2A,B), indicating that M6 is not required for trafficking of Aka to the plasma membrane, but for accumulation of Aka at TCJs. Furthermore, immunoblots revealed that the total amounts of full-length Aka protein were reduced to 69.4 ± 8.9% (n=2) in *M6^W186^\** embryos and to 38.3 ± 1.6% (n=2) in *M6^MB02608^* embryos compared to controls (Fig. 2Q). Together these results indicate that M6 is required for the assembly of tricellular, but not bicellular septate junctions in embryonic epithelia, and suggest that the stability of Aka protein depends on M6.

### M6 acts in a cell-autonomous fashion

To corroborate that the defects in TCJ formation were caused by the absence of M6, we generated a transgene in which the genomic *M6* locus with a C-terminal GFP tag was placed downstream of UAS sequences for Gal4-driven expression (UAS-M6::GFP; Fig. 1A). When expressed in epidermal stripes in *M6^MB02608^* embryos, M6::GFP localized to lateral membranes and accumulated at TCJs (Fig. 2O”,P”). Moreover, M6::GFP rescued the localization of Aka (Fig. 2O) and Gli (Fig. 2P) to vertices in the M6::GFP-expressing stripes, whereas Aka and Gli were mislocalized around lateral membranes in neighboring cells. Thus, M6 is required in a cell-autonomous fashion for the localization of Aka and Gli to TCJs in epidermal cells.

### Aka and M6 act in a mutually dependent manner upstream of Gli to promote TCJ assembly

To understand how Aka, Gli and M6 cooperate in TCJ assembly, we analyzed by immunostaining the distribution of the three TCJ components in loss-of-function mutants of each gene (*aka^L200^, M6^MB02608^, Gli^F156^*; Fig. 3A-D). While Aka, Gli and GFP::M6 accumulated at TCJs in epidermal cells of stage 15 wild-type embryos (Fig. 3A), accumulation of Aka and Gli at TCJs was lost in the absence of M6, as described above (Fig. 2K,M, Fig. 3C; Fig. S2). Similarly, in the absence of Aka, both GFP::M6 and Gli failed to accumulate at vertices and instead were spread around the cell perimeter (Fig. 3B). By contrast, in the absence of Gli both Aka and GFP::M6 accumulated at vertices, comparable to their distribution in wild-type embryos (Fig. 3D). Thus, Aka and M6 depend on each other, but not on Gli, for their localization to TCJs, whereas Gli depends on both Aka and M6 (Fig. 3F). Together these findings indicate that Aka and M6 synergize to promote TCJ assembly by recruiting Gli to vertices.

**Figure 3.**
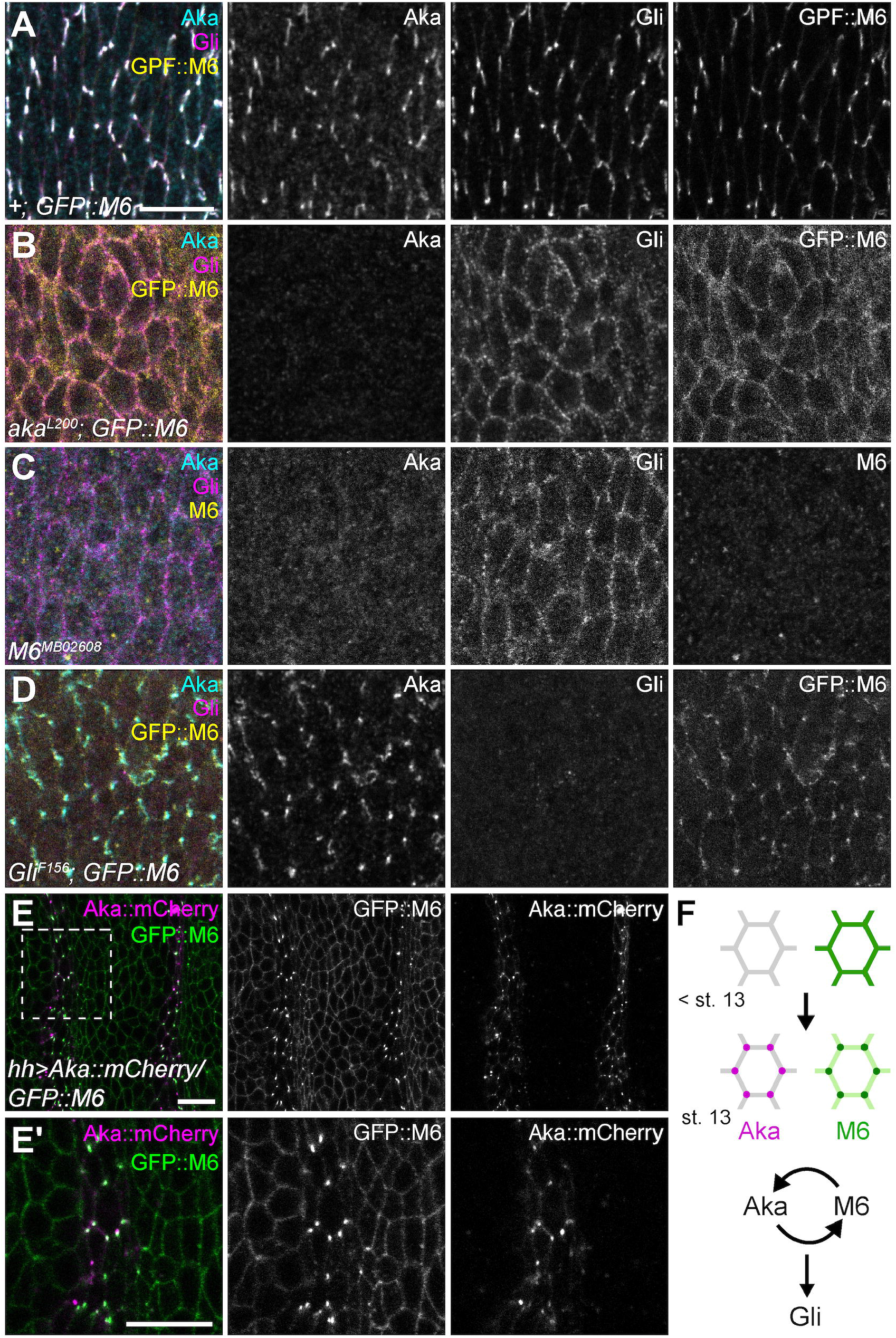
Aka and M6 act in a mutually dependent manner upstream of Gli to promote TCJ assembly. (A-D) Embryos (stage 15) of the indicated genotypes were stained for Aka and Gli. M6 was detected by GFP::M6 fluorescence (A,B,D) or anti-M6 immunostaining (C). In *+; GFP::M6* control embryos (A) Aka, Gli, and GFP::M6 accumulate at TCJs, whereas in *aka^L200^* mutants (B) Gli and GFP::M6 are mislocalized. Similarly, in *M6^MB02608^* mutants (C) Aka and Gli are mislocalized. By contrast, in *Gli^F156^* mutants (D), Aka and GFP::M6 accumulate at TCJs. (E,E’) Premature expression of Aka::mCherry in epidermal stripes of stage 12 *GFP::M6* embryos (driven by *hh*-Gal4) is sufficient for accumulation of GFP::M6 at TCJs in the Aka::mCherry-expressing cells, whereas GFP::M6 is distributed around the lateral membrane in neighboring cells. (E’) shows close-up of region marked by rectangle in (E). (F) Scheme summarizing the dynamics of Aka and M6 localization at TCJs (top) and relationships between requirements of Aka, M6 and Gli (bottom). See text for details. Scale bars: 10 μm.

### Aka, M6, and Gli accumulate simultaneously at TCJs

Having established that Aka and M6 act in an interdependent manner upstream of Gli, we analyzed the kinetics of TCJ formation to address whether the proteins assemble sequentially or simultaneously. Zygotic expression of M6 and Gli starts during germband elongation (stage 10), approximately six hours before the onset of Aka expression, which sets in after completion of germband retraction (stage 13), as revealed by RNA-Seq [19]; FlyBase), mRNA *in situ* hybridization and immunostainings [6, 16]. Interestingly, at early stage 13 GFP::M6 and Gli were distributed around the perimeter of epidermal cells (Fig. S1J), while at late stage 13, GFP::M6 and Gli began to accumulate at vertices, first observable in segmental patches in the dorsal epidermis (Fig. S1K) and subsequently spreading throughout the epidermis (Fig. S1L). This change in localization of M6 and Gli coincided with the onset of expression of Aka, which accumulated at vertices together with GFP::M6 and Gli (Fig. S1K). Intriguingly, all vertices displaying accumulation of Aka also contained GFP::M6 and Gli at comparable intensities between different vertices (Fig. S1K,L), suggesting that Aka, M6, and Gli assemble simultaneously at TCJs. Live imaging revealed that GFP::M6 rapidly accumulates at vertices, first appearing in puncta that subsequently extend basally along vertices (Fig. S1M,N; Movie S1), suggesting that TCJ assembly involves dynamic redistribution of M6 protein between bicellular contacts and vertices.

### Aka is sufficient to direct M6 to vertices in the absence of septate junctions

As the accumulation of M6 and Gli at TCJs coincided with the appearance of Aka at stage 13, we asked whether premature expression of Aka could be sufficient to direct GFP::M6 to vertices prior to stage 13. Strikingly, expression of Aka::mCherry in epidermal stripes of stage 12 embryos indeed led to accumulation of GFP::M6 at TCJs in the Aka::mCherry-expressing cells, whereas GFP::M6 was distributed around the cell perimeter in the neighboring cells (Fig. 3E). To test whether this effect was dependent on the presence of bicellular SJs, we expressed M6::GFP and Aka in stage 11 embryos, which have not yet formed SJs ([20]; Fig. S3). When M6::GFP or Aka::mCherry were expressed individually, M6::GFP was distributed around the lateral membrane (Fig. S3A,E) and Aka::mCherry was visible in cytoplasmic puncta (Fig. S3B,E). Strikingly, however, when M6::GFP and Aka::mCherry were co-expressed, the two proteins accumulated together at vertices (Fig. S3C,D,E). These results indicate that Aka is sufficient to direct M6 to vertices in the absence of SJs.

### Aka and M6 are mutually dependent for their low mobility at vertices

To understand the mode of action of Aka and M6 we analyzed the mobility of endogenously tagged Aka::GFP and GFP::M6 at bicellular junctions (BCJs) and TCJs, respectively, using Fluorescence Recovery After Photobleaching (FRAP) experiments (Fig. 4). At epidermal TCJs, Aka::GFP (Fig. 4A-C) and GFP::M6 (Fig. 4G-I) showed slow recovery with small mobile fractions (Aka::GFP: 23%, n=12; GFP::M6: 10%, n=11), comparable to SJ proteins, such as Nrg::YFP (18%, n=8; Fig. S4), which form immobile complexes [21]. However, unlike Nrg::YFP or the cytoplasmic SJ-associated protein Discs-large (Dlg::mTagRFP; Fig. S4), Aka::GFP (Fig. 4A-C) and GFP::M6 (Fig. 4G-I) displayed substantially larger mobile fractions at BCJs than at TCJs (Aka::GFP: 46%, n=12; GFP::M6: 35%, n=11), suggesting that BCJs and TCJs comprise distinct pools of Aka and M6 proteins. Interestingly, the low mobility of GFP::M6 at TCJs disappeared in the absence of Aka, and mobility at TCJs and BCJs became indistinguishable (Fig. 4J-L), suggesting that without Aka, M6 becomes equally mobile around the plasma membrane. Correspondingly, in the absence of M6, Aka::GFP showed dramatically increased mobility both at TCJs and BCJs, with mobile fractions larger than in wild-type embryos (Fig. 4D-F). These findings indicate that Aka and M6 are mutually dependent for their accumulation and stabilization in a low-mobility pool at cell vertices.

**Figure 4.**
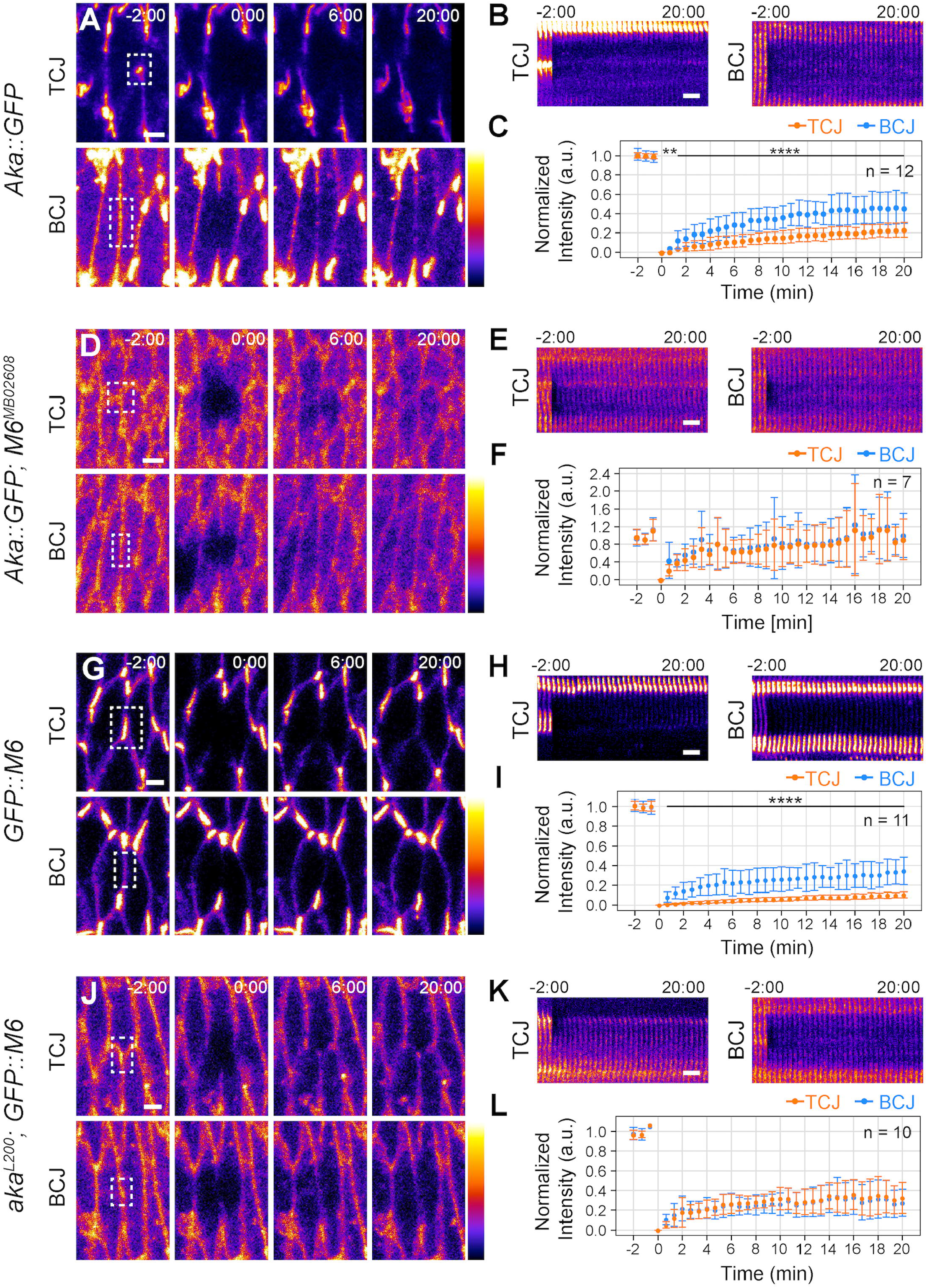
Aka and M6 are mutually dependent for their low mobility at vertices. (A-L) FRAP was performed in epidermis of stage 15 embryos of the genotypes indicated. (A,D,G,J) show stills of representative movies. Bleached regions at TCJs (top) or BCJs (bottom) are indicated by boxes. (B,E,H,K) show kymographs of bleached regions. (C,F,I,L) show fluorescence recovery over time. Data is represented as mean +/- S.D. Number of embryos analyzed (n) per genotype is indicated. Wilcoxon rank-sum test, **: p < 0.01, ****: p < 0.0001. Note that Aka::GFP shows a smaller mobile fraction at TCJs compared to BCJs in wild-type control (A-C), but becomes highly and equally mobile at BCJs and TCJs in the absence of M6 (D-F). Similarly, GFP::M6 shows a smaller mobile fraction at TCJs compared to BCJs in controls (G-I), but the low-mobility TCJ fraction is lost and GFP::M6 becomes equally mobile at TCJs and BCJs in the absence of Aka (J-L). Kymographs (B,E,H,K) indicate that lateral diffusion contributes little to recovery of Aka::GFP and GFP::M6 on the observed time scale. See also Figure S4. Scale bars: 2 μm.

Taken together, we showed that the transmembrane proteins M6 and Aka assemble at TCJs in a mutually dependent fashion and are required jointly for TCJ localization of Gli. Conversely, Gli is not required to bring Aka or M6 to TCJs, suggesting that Aka and M6 initiate TCJ assembly, while Gli is recruited downstream and not required for the initial assembly of TCJ complexes. Aka acts instructively, as its premature expression is sufficient to direct M6 to cell vertices in the absence of septate junctions. Conversely, M6, which is expressed before Aka, appears to be required permissively to maintain Aka localized at TCJs, but not for trafficking of Aka to the cell surface. Finally, we demonstrate that cell vertices comprise a distinct low-mobility pool of TCJ proteins, while the proteins display substantially higher turnover at bicellular junctions. Aka and M6 are interdependent to maintain their accumulation and low mobility at TCJs, suggesting that they are part of a complex that forms or becomes stabilized selectively at vertices. Thus, we delineate a pathway of TCJ assembly in which Aka and M6 promote TCJ formation through synergistic interactions that establish a distinct plasma membrane domain at cell vertices.

What could direct the formation of this membrane domain? Our findings suggest that the unique geometry of epithelial cell vertices provides specific cues that promote the assembly of TCJ complexes. The apposition of three convex plasma membranes towards the extracellular TCJ space could be sensed by Aka’s large triple-repeat extracellular domain, which we showed mediates cell adhesion [6]. Conversely, on the cytosolic side, the convex membrane at the vertex is likely to be enriched in specific lipids, including cholesterol [22]. Interestingly, the vertebrate M6 homolog GPM6a was shown to associate with cholesterol-rich, negatively curved membranes and to induce the formation of membrane protrusions, such as neurites, filopodia, and dendritic spines in neurons in a cholesterol-dependent manner [11, 23-25]. In a similar fashion, membrane curvature at vertices could promote accumulation of M6 protein.

M6/PLP proteins contain four transmembrane domains that share structural similarity with tetraspanin family proteins [26]. Moreover, like tetraspanins, which bind cholesterol in a palmitoylation-dependent manner [27], *Drosophila* M6 and mouse M6a are palmitoylated [10, 28], and mouse M6a segregates into cholesterol-enriched membrane domains in a palmitoylation-dependent fashion [10]. Tetraspanins associate with each other and with various transmembrane proteins, as well as with membrane regions enriched in cholesterol and glycosphingolipids, forming tetraspanin-enriched microdomains [29]. Resembling the role of tetraspanins, mammalian M6a and M6b are required for correct plasma membrane localization of transmembrane proteins including the μ-opioid receptor and the serotonin transporter [30, 31]. It will be interesting to test whether M6 acts in an analogous fashion in TCJ assembly by promoting the maintenance of Aka at a vertex-specific membrane nanodomain, and whether palmitoylation of M6 and cholesterol are required. Intriguingly, TCJ localization of the transmembrane protein Angulin-1 in mammalian cells was shown to depend on palmitoylation of the Angulin-1 cytosolic domain and on plasma membrane cholesterol, suggesting that Angulin-1 segregates into a membrane domain defined by its specific lipid composition [32]. Thus, although TCJs are comprised of different proteins in vertebrates and invertebrates, common principles appear to mediate the vertex-specific assembly of TCJ complexes.

## Materials and Methods

### *Drosophila* strains and genetics

The following *Drosophila* stocks are described in FlyBase, unless noted otherwise: *aka^L200^, UAS-Aka, UAS-Aka::mCherry, Gli^F156^* [6], *M6^W186*^, M6^NC^, M6^G152E^* [8], *M6^MB02608^* [15], *Dlg::mTagRFP* [33], *Gli::YFP^CPTI-002805^, Nrg::YFPC^PTI001714^* [34], *UAS-palm-neonGreen* [35], *CyO Dfd-GMR-nvYFP, TM6B Dfd-GMR-nvYFP* [36], *en-Gal4, UAS-mCherry-nls, Df(3L)BSC419. Aka::GFP* carries a CRISPR-mediated insertion of superfolder-GFP (sfGFP [37]) into the cytoplasmic domain of Aka (23 amino acids from the C-terminus) in the *aka* locus (gift from Maite Vidal-Quadras and Anne Uv, University of Gothenburg, Sweden).

### Molecular biology

UAS-M6 was generated as follows: A genomic fragment comprising the *M6* coding region (including the 5’- and 3’-UTRs of isoforms C, D, E, F, G) was amplified from genomic DNA of *y^1^ w^1118^* flies and ligated into pCR-BluntII-Topo using the Blunt TOPO Cloning kit (Thermo Fisher), resulting in pCR-BluntII-TOPO-M6-fl. The *M6* insert was excised with NotI and KpnI and ligated into pUASt-attB [38].

UAS-M6::GFP was generated as follows: A 3’ fragment of the *M6* sequence from the NheI site to the last codon preceding the stop codon was amplified by PCR from pCR-BluntII-TOPO-M6-fl using oligonucleotides with overlaps to the upstream *M6* sequence and to the sfGFP coding sequence. The sfGFP coding sequence, including a stop codon, was amplified from Aka-TY1-TEV-sfGFP-FLAG [6] using primers with overlaps to *M6* and the pCR-BluntII-TOPO vector. pCR-BluntII-TOPO-M6-fl (see above) was cut with NheI and KpnI, and the 3’- *M6* fragment and the sfGFP fragment were inserted using In-fusion cloning (homologous recombination; TaKaRa). The resulting M6-sfGFP insert was excised using NotI and KpnI and ligated into pUASt-attB [38]. The resulting UAS-M6::GFP construct is expected to produce C-terminally GFP-tagged versions of M6 isoforms C, D, E, and F, but not of the C-terminally truncated RG isoform and of transcript RB, which is expressed from a downstream promotor (Fig. S1A).

The pUASt-attB-M6 and pUASt-attB-M6-sfGFP constructs were inserted into the attP40 and attP2 landing sites, respectively, using PhiC31-mediated site-specific integration [38].

### Antibodies and Immunostaining

Embryos were fixed in 4% formaldehyde in PBS/heptane for 20 min and devitellinized by shaking in methanol/heptane. Antibodies were mouse anti-Gli 1F6.3 (1:500; [7]), rabbit anti-Aka (1:500; [6]), chicken anti-mCherry (1:1000; Novus Biologicals), rabbit anti-Mcr (1:1000; [39]), mouse anti-Cora C615.16 (1:200; [18]), mouse anti-Fas3 7G10 (1:50; [40]) and rabbit anti-Nrx-IV (1:100; [17]). Goat secondary antibodies were conjugated with Alexa 488, DyeLight 550, Alexa 568 or Alexa 647 (Life Technologies). The anti-M6 antiserum was generated by immunizing guinea pigs (Eurogentec) with the peptides RRNSYRSDHSLDRYT and NLNELEYSATSKDRF (corresponding to aa 102-116 and 354-368 in M6 isoform F) and was used at a dilution of 1:500 for immunostainings.

### Immunoblots

25 embryos (stage 15) were homogenized in 21μl lysis buffer (50mM Tris/HCl, 150mM KCl, 5mM MgCl_2_, 1x Halt protease inhibitor cocktail (pH 8.0; Thermo Fisher)) on ice. After adding 7μl Lämmli buffer (250mM Tris, 8% SDS, 40% glycerol, 20% beta-mercaptoethanol, 0.2% bromophenol blue; pH 6.8) samples were boiled at 95°C for 5 min. 12μl of sample were loaded on 4%-20% Mini-PROTEAN TGX gels (Bio-Rad) and blotted on PVDF membranes (pore size 0.45μm; Amersham). Primary antibodies (rabbit anti-Aka (1:5000; [6], mouse anti-alpha-Tubulin AA4.3-c (1:5000; DSHB)) were detected using HRP-conjugated secondary antibodies (1:10000; Thermo Fisher) and ECL Prime kit (Amersham). Raw images of immunoblots were processed using scikit-image (v0.16.2). Lanes of equal width were selected and the intensity profiles calculated as the rowsum of pixel intensity values. Background (30%-quantile of all the rowsums) was subtracted from the raw values. Intensities for all protein bands were calculated and corrected for total amount of protein by the intensity of the alpha-Tubulin band (loading control).

### Dextran injections

Rhodamine-labeled dextran (10 kDa; Sigma) was injected into stage 16 control (*y^1^ w^1118^*) and homozygous *M6^MB02608^* embryos (identified by the absence of the Dfd-YFP-expressing balancer chromosome) as described [41]. Embryos were imaged 20 minutes after injection and the distribution of dextran was analyzed.

### Microscopy and image analysis

Imaging was performed on a Leica SP8 confocal microscope with 40x/1.3 NA and 63x/1.4 NA objectives. For live imaging, dechorionated embryos were mounted on glue-coated coverslips (0.17mm, grade #1.5) and covered with Voltalef 10S oil (VWR). Movies of GFP::M6 accumulation were acquired using a 40x/1.3 NA oil immersion objective and the SP8 resonant scanner (line accumulation of 2, frame averaging of 6) at 30s time intervals. Images were processed using OMERO.web (5.5.1), OMERO.figure (4.2.0), and FIJI (ImageJ) (1.52p [42]).

### Quantification of protein accumulation at tricellular junctions

Embryos (stage 15) were staged according to gut morphology, aligned dorsally and mounted as described above. Z-stacks with 10 slices were acquired with the following settings: 640 x 640px image size, 4x line accumulation, 400Hz, 40x/1.3 NA objective, HyD detectors in counting mode and 1.5% laser intensity of the 488nm laser. For quantification of protein enrichment at vertices, average projections of the three most apical slices showing junctional signal in each stack were analyzed. Mean fluorescence was measured at tricellular contacts (TCJs), bicellular contacts (BCJs) and inside the cytoplasm using equally sized ROIs (radius 20px/1.52μm). Enrichment was calculated as

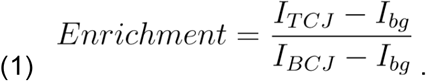

At least 20 TCJs, 40 BCJs and 20 background measurements were analyzed per image. Vertices bordered by more than three cells were not included in the measurements. Intensities in each category (TCJ, BCJ, cytoplasm) were averaged for each image and one enrichment value per image was calculated. The sample size (n) indicated in the figures states the number of embryos analyzed.

### Measurements of tracheal tube length

The tracheal lumen in embryos (stage 16) was visualized by staining for chitin using Alexa-Fluor-SNAP-tagged chitin-binding domain from *Bacillus circulans* chitinase A1 prepared as described in [43]. To measure the length of the tracheal dorsal trunk, confocal z-stacks covering the entire volume of the dorsal trunk were acquired. The length of the dorsal trunk was measured from the second to the eighth dorsal trunk metamere in three dimensions using the filament tool in Imaris (v 9.0.0) with manual positioning of the filament.

### Fluorescence recovery after photobleaching experiments

Embryos (stage 15) were staged according to gut morphology, and mounted on glue-coated coverslips with the dorsal side facing the coverslip. Image acquisition was performed using the Live Data Mode (Leica LAS X) with the following settings: 480px x 480px image size, 4x line accumulation, 1000Hz, 40s time intervals, 40x/1.3 NA objective, and HyD detectors in counting mode. Jobs in the Live Data Mode were set to the following conditions: (i) Prebleach series with three time frames, followed by (ii) a 30s pause to draw regions to be bleached, (iii) bleach step using 10 repetitions without pause with 100% laser intensity at 488nm and 552nm at 200Hz, and (iv) post-bleach series with 31 time frames. Z-drift was corrected manually during acquisition. Average projections of FRAP time series were generated using the three most apical slices showing junctional signal. XY-drift in the series was corrected using the Template matching plugin [44] in ImageJ. Images were cropped to exclude any black pixels originating from XY-drift correction. Regions of Interest (ROIs) were drawn around bleached regions and mean intensity was recorded at every time point. Normalized intensity was calculated using the following formulas:

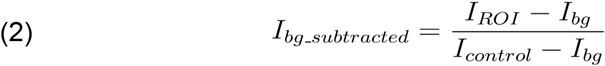

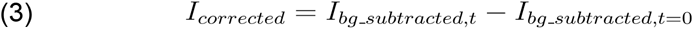

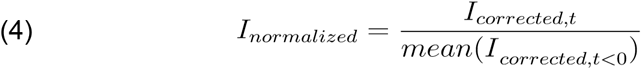

with *I_ROI_* representing the measured intensity inside the ROI, *I_control_* representing the bleach control (described below), *I_bg_* representing the background (described below), *I_bg subtracted,t=0_* representing the background-subtracted intensity immediately after bleaching, and *I_corrected,t<0_* representing the background-subtracted mean intensity for all time points before bleaching. The bleach control was calculated as the mean intensity of the brightest 5% of pixels in an image (selected pixels correspond to TCJs). The background was calculated as the mean intensity of the dimmest 40% of pixels in an image (selected pixels correspond to the cytoplasm). Correction and normalization were performed individually for each ROI using the image-wide background and bleach control. Curves were fitted to the normalized data using R and the following formula with A = 1 and τ = 1 as starting values:

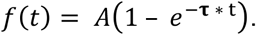

For displaying FRAP curves, each time point is shown as the mean +/- standard deviation. Wilcoxon rank-sum test was performed for the values at each time point. Kymographs were generated using a self-written ImageJ script. A rectangular 12px x 130px (0.91μm × 9.85μm) selection was duplicated for each time point and assembled horizontally to create the kymograph.

### Statistics

For phenotypic analyses, sample size (n) was not predetermined using statistical methods, but was assessed by taking into account the variability of a given phenotype, determined by the standard deviation. Experiments were considered independent if the specimens analyzed were derived from different parental crosses. During experiments investigators were not blinded to allocation. Data was checked for normal distribution using the Shapiro-Wilk test. If the groups to be compared were normally distributed, Welch’s t-test for unequal variances (R standard package) was used. If one group was not normally distributed, the Wilcoxon ranksum test was used. P-values were corrected for multiple testing using the Bonferroni-Holm method [45].

## Acknowledgements

We thank Wilko Backer for expert technical help, Roland Le Borgne, Maite Vidal-Quadras and Anne Uv for sharing unpublished data and for comments on the manuscript, and Yohanns Bellaiche, Fernanda Ceriani, Maite Vidal-Quadras, Anne Uv and Tian Xu for providing fly stocks and reagents. Work in SL’s laboratory was supported by the Deutsche Forschungsgemeinschaft (SFB1348 “Dynamic Cellular Interfaces”; SFB1009 “Breaking Barriers”), the “Cells-in-Motion” Cluster of Excellence (EXC 1003-CiM) and the University of Münster.

---

**Figure S1.**
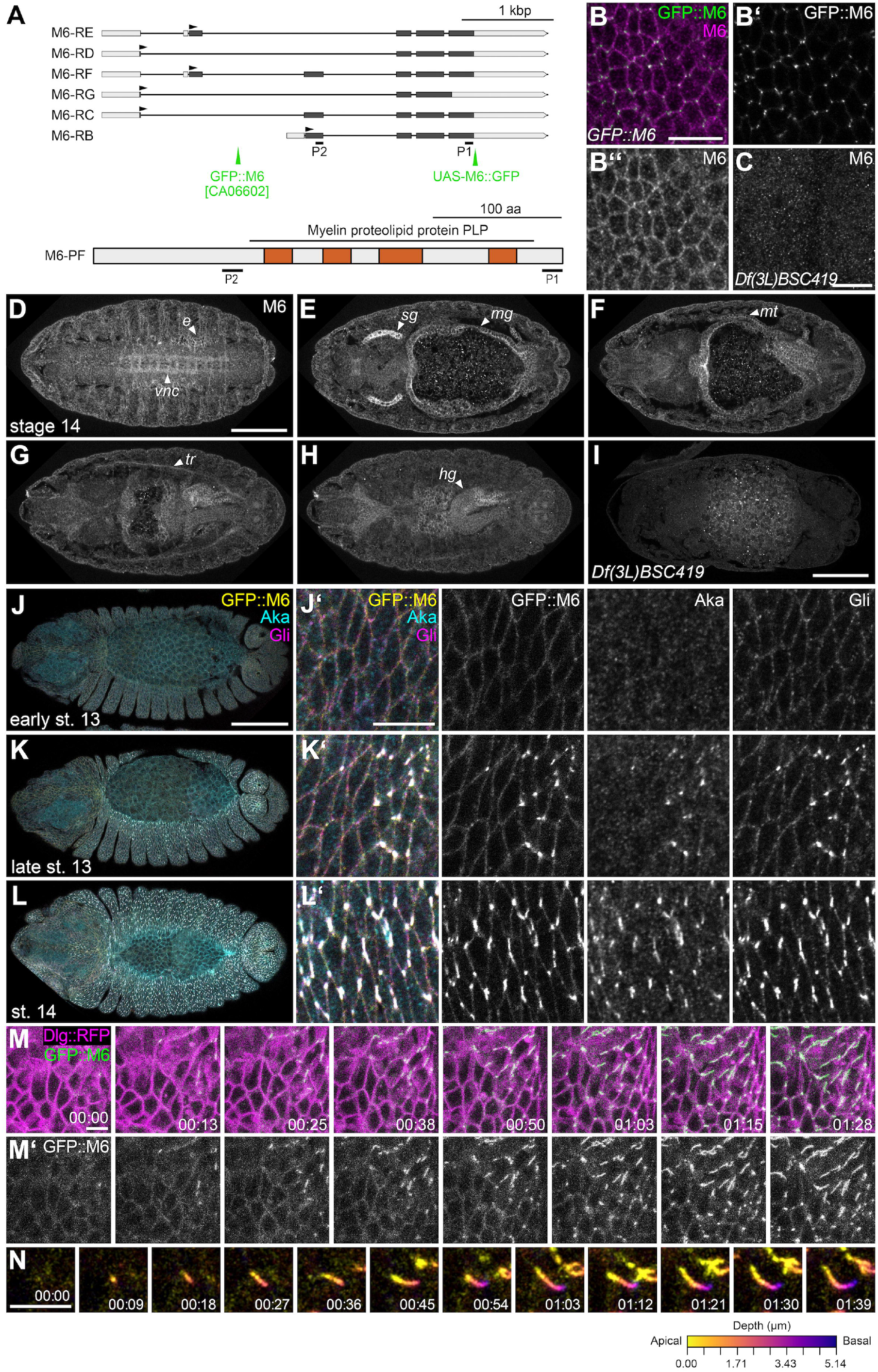
M6 is detectable in embryonic epithelia and accumulates simultaneously with Aka and Gli at TCJs. (A) Schematic representation of *M6* transcript isoforms (top; according to FlyBase). Exons are shown as boxes with coding portions in black and non-coding portions in grey. Black arrowheads mark positions of translation start sites. Green arrowheads mark the positions of GFP in the CA06602 protein trap and in the UAS-M6::GFP construct. The anti-M6 antiserum was raised against the peptides indicated as P1 and P2. The domain organization of M6 protein (isoform F) is shown at the bottom. Predicted transmembrane domains are shown in orange. The region with homology to vertebrate myelin proteolipid proteins is indicated. Isoforms B, C, D, E and F share the four transmembrane domains and a common C-terminus, while isoform G is predicted to lack the fourth transmembrane domain. (B,C) Epidermis in stage 15 *GFP::M6* embryo (B-B’’) stained with anti-M6 antiserum (B’’; magenta in merge). GFP::M6 was detected as GFP fluorescence (B’; green in merge). Note that anti-M6 staining overlaps with GFP::M6 signals at TCJs, but additionally labels lateral cell membranes. (C) Absence of junctional signal in epidermis of stage 15 *Df(3L)BSC419* embryo stained with anti-M6 indicates specificity of the anti-M6 antiserum. (D-I) Confocal sections of stage 14 wild-type embryo stained with anti-M6 antiserum. Different sections of the same embryo are shown. M6 staining is visible in the epidermis (*e*; D), ventral nerve cord (*vnc*; D), salivary glands (*sg*; E), midgut (*mg*; E), Malpighian tubules (*mt*; F), tracheae (*tr*, G), and the hindgut (*hg*; H). Absence of staining in *Df(3L)BSC419* homozygous embryo (I) indicates specificity of the anti-M6 antiserum. (J-L) Dorsal views of embryos (J,K,L; z-pro) and close-ups of dorsal epidermis (J’,K’,L’) in *GFP::M6* embryos at early stage 13 (J), late stage 13 (K) and stage 14 (L) stained for Aka (cyan) and Gli (magenta). GFP::M6 was detected as GFP fluorescence (yellow). Note that GFP::M6 and Gli are distributed around lateral cell membranes at early stage 13 (J’) and begin to accumulate at TCJs together with Aka at late stage 13, first in patches in the dorsal epidermis (K’) and throughout the epidermis at stage 14 (L’). (M) Still series (maximum-intensity z-projections) from time-lapse movie (Movie S1) of late stage 13 embryo expressing GFP::M6 (green) and Dlg::mTagRFP (Dlg::RFP; magenta). A close-up of dorsal epidermis is shown. Anterior is to the left, dorsal is up. Time is indicated (hh:mm). GFP::M6 is initially distributed around lateral membranes and begins to accumulate at vertices at 13 minutes. Accumulation at vertices spreads from dorsal to ventral epidermal cells. Note that GFP::M6 signals initially appear as puncta and that signals extend along vertices over time. See also movie S1. (N) Close-ups (maximum-intensity z-projections) of GFP::M6 at a single vertex. Signals were color-coded for depth using the heat-map scale shown to the lower left. Note that GFP::M6 accumulation extends along the vertex in an apical to basal direction. Scale bars: (B,C,J’,K’,L’,M), 10 μm; (D-H,I,J-L), 100 μm; (N), 5 μm.

**Figure S2.**
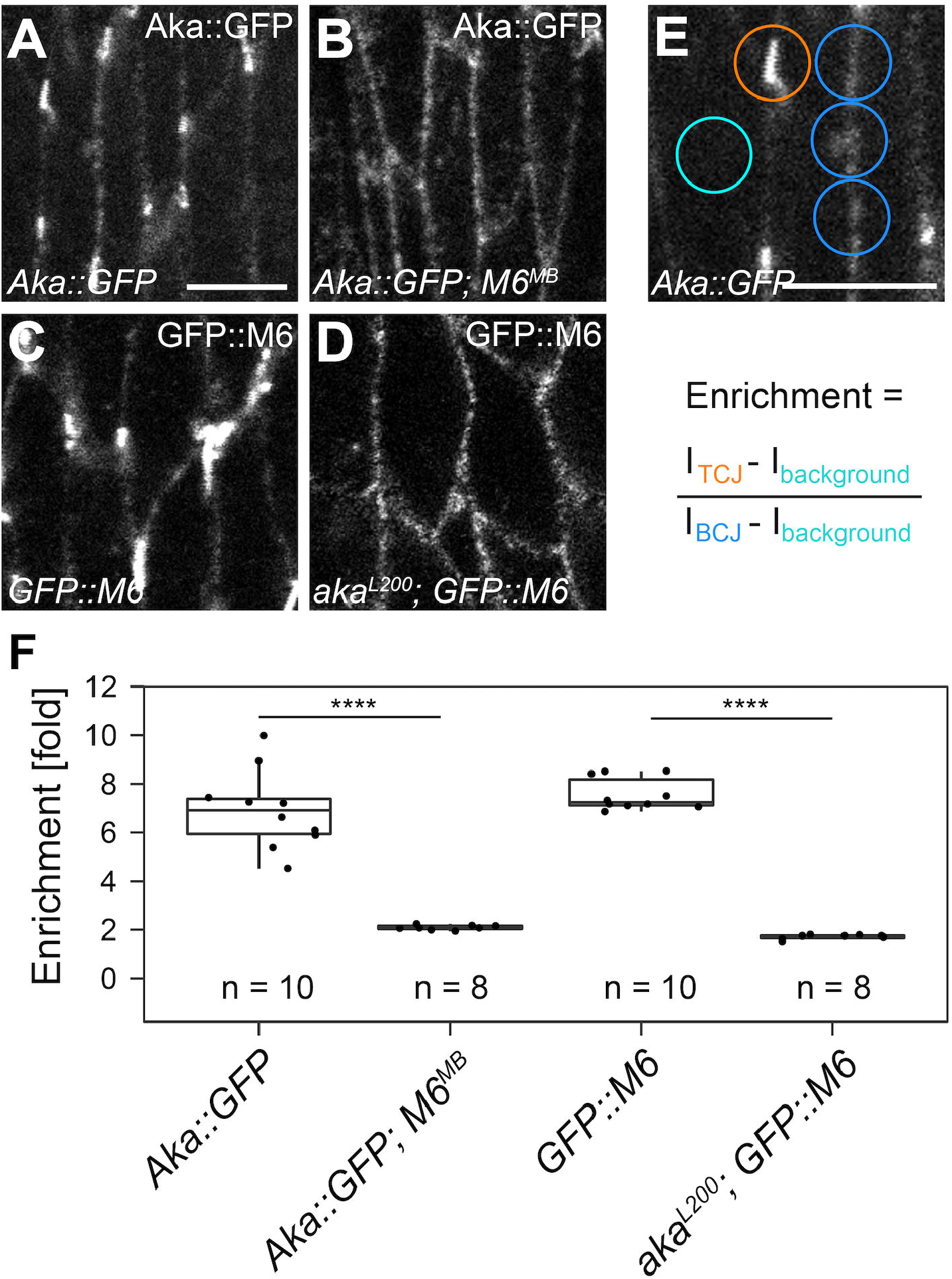
Aka and M6 localize to tricellular junctions in a mutually dependent fashion. (A-E) En-face view of epidermis in living embryos (stage 15) expressing Aka::GFP in wildtype (A) or *M6^MB02608^* mutant (B) background, or expressing GFP::M6 in wild-type (C) or *aka^L200^* mutant (D) background. Note that Aka::GFP and GFP::M6 accumulate at vertices in wild-type embryos, but are distributed around lateral membranes in *M6^MB02608^* or *aka^L200^* embryos, respectively. (E) Quantification of protein accumulation at TCJs. Equally sized ROIs were used to measure fluorescence intensities at TCJs (orange), BCJs (blue) or inside the cytoplasm (cyan). Enrichment was calculated as the ratio of background-subtracted intensity at TCJs to background-subtracted intensity at BCJs. (F) Box plot showing enrichment (fold) of fluorescence at TCJs in the indicated genotypes. Note that Aka::GFP and GFP::M6 accumulate at TCJs in a mutually dependent manner. Number of embryos analyzed (n) per genotype is indicated. Wilcoxon rank-sum test, ****: p < 0.0001. Scale bars: 5 μm.

**Figure S3.**
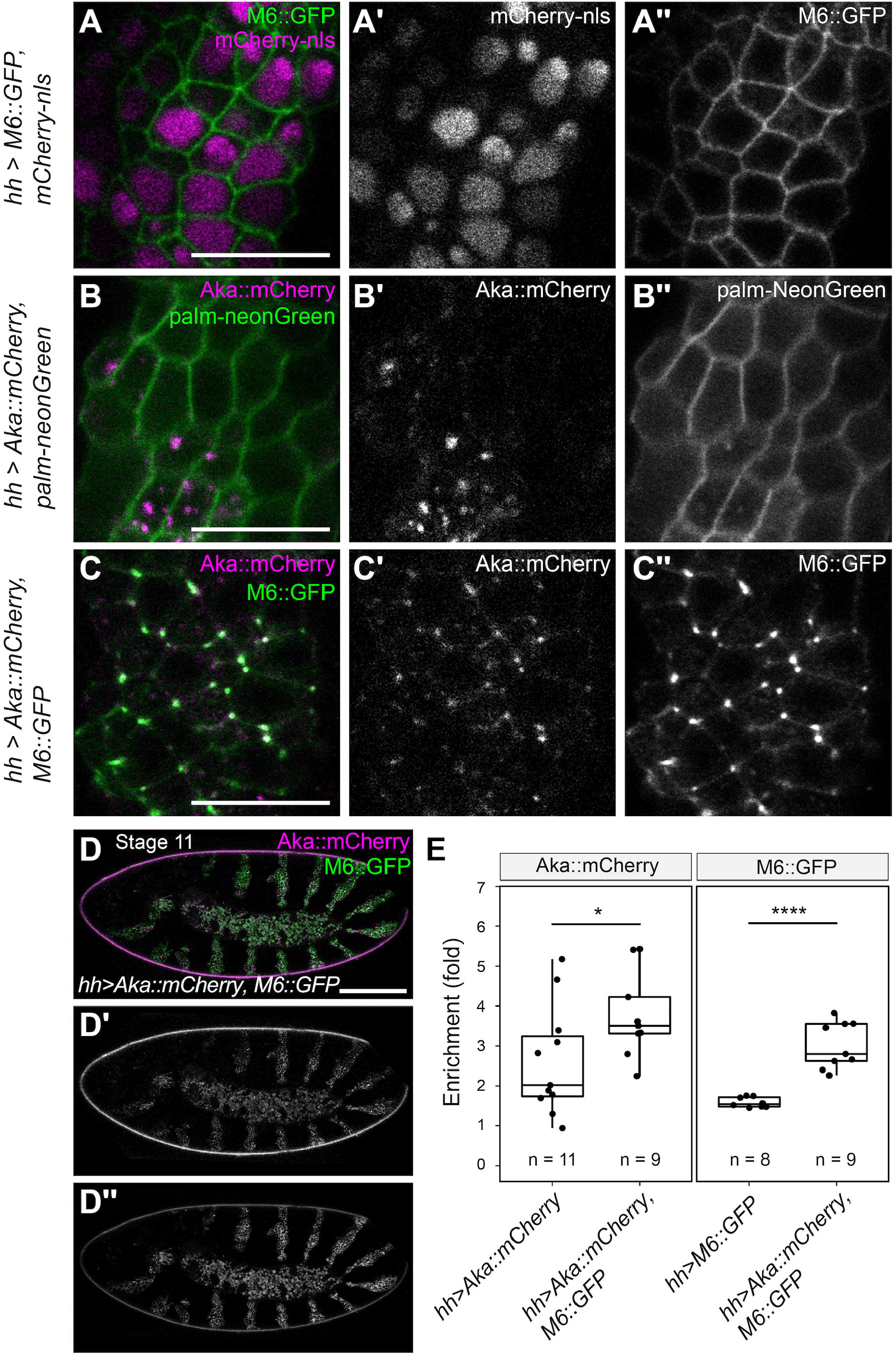
Aka is sufficient to direct M6 to vertices in the absence of septate junctions. (A-D) Confocal sections of epidermis in living embryos (stage 11) expressing M6::GFP and mCherry-nls (A), Aka::mCherry and palm-neonGreen (B), or Aka::mCherry and M6::GFP (C,D) under the control of *hh*-Gal4 in epidermal stripes. (D-D’’) shows overview of stage 11 embryo expressing Aka::mCherry and M6::GFP. Note that co-expression of Aka::mCherry and GFP::M6 leads to accumulation of both proteins at vertices (C), while GFP::M6 alone is distributed around lateral membranes (A’’) and Aka::mCherry localizes to intracellular puncta (B’). (E) Box plot showing enrichment (fold) of fluorescence at TCJs in the indicated genotypes. Number of embryos analyzed (n) per genotype is indicated. Wilcoxon rank-sum test, *: p < 0.05; ****: p < 0.0001. Scale bars: (A-C), 5 μm; (D-D’’) 100 μm.

**Figure S4.**
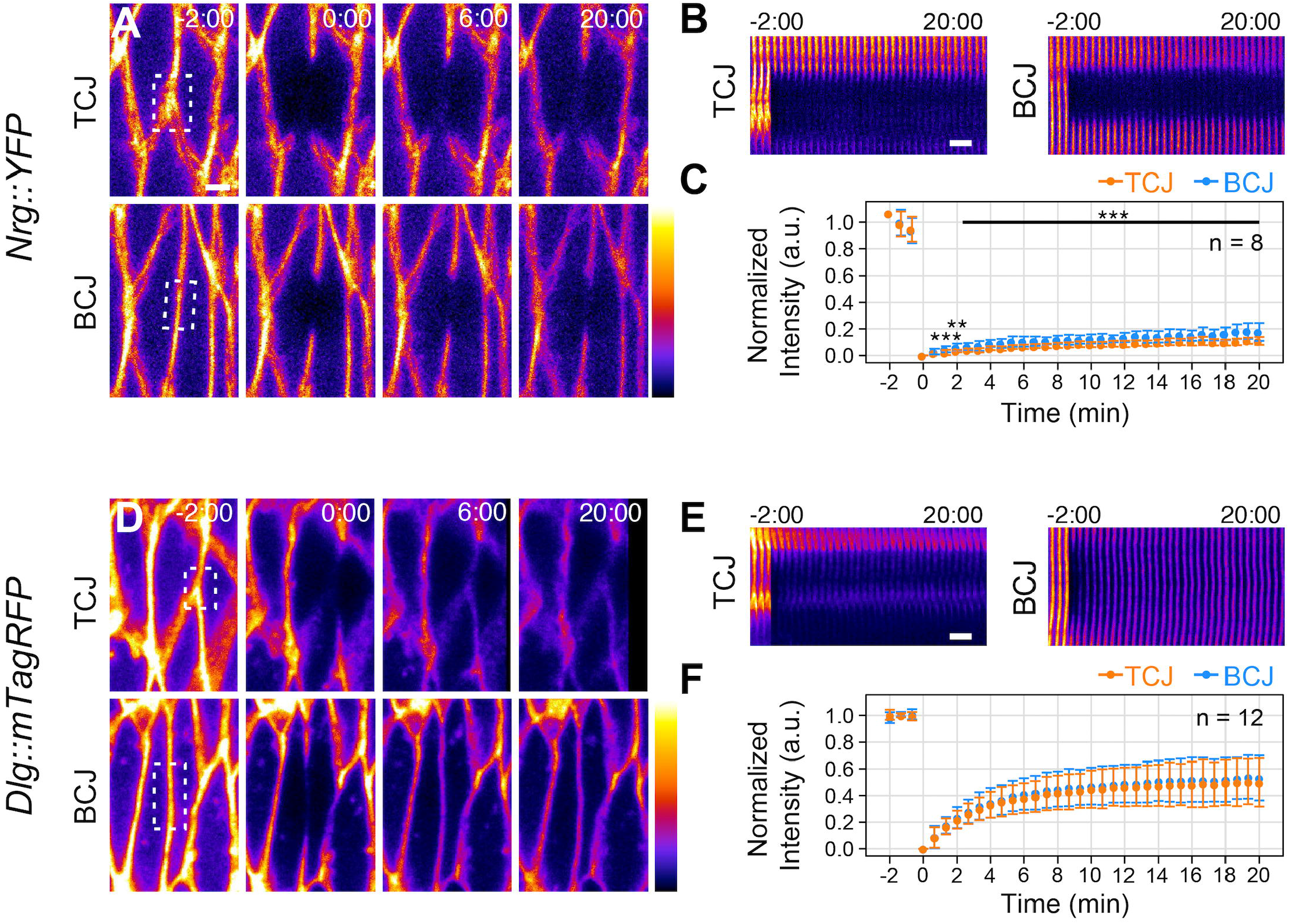
Mobility of septate junction proteins Nrg and Dlg is similar at bicellular and tricellular junctions. (A-F) FRAP was performed in the epidermis of stage 15 embryos of the genotypes indicated to the left. (A,D) show still series of representative movies. Bleached regions at TCJs (top panels) or BCJs (bottom panels) are indicated by dashed boxes. (B,E) show kymographs of bleached regions. (C,F) show fluorescence recovery over time. Data is represented as mean +/- S.D. Number of embryos analyzed (n) per genotype is indicated. Wilcoxon rank-sum test, **: p < 0.01, ***: p < 0.001. Note that the SJ transmembrane protein Nrg::YFP (C) displays slow recovery with a low mobile fraction below 20%, whereas the SJ-associated cytoplasmic protein Dlg::mTagRFP displays fast recovery with a high mobile fraction of 50% (F), consistent with [21]. However, unlike the TCJ proteins Aka::GFP and GFP::M6 (Fig. 4), both Nrg::YFP and Dlg::mTagRFP show similar mobility at BCJs and TCJs. Kymographs (B,E) indicate that lateral diffusion contributes little to recovery of Nrg::YFP (B), but dominates recovery of Dlg::mTagRFP at BCJs (E). Scale bars: 2 μm.

## Supplemental movies

**Movie S1**

**Time-lapse movie of GFP::M6 accumulation at TCJs in a stage 13 embryo.**

Maximum intensity projection of lateral epidermis in a *Dlg::mTagRFP;; GFP::M6* embryo (stage 13). Anterior is to the left, dorsal is up. GFP::M6 (green in merge) is initially distributed around lateral membranes and begins to accumulate at vertices in segmental patches in the dorsal epidermis. Note that GFP::M6 accumulation at vertices spreads from dorsal to ventral in the epidermis, and that GFP::M6 signals extend in an apical to basal direction along each vertex (see also Figure S1). Images were acquired at 30s intervals. Time is indicated (hh:mm:ss).

Scale bar: 10 μm.

## Notes

### Competing Interest Statement

The authors have declared no competing interest.

